# The Serine Protease HtrA2 mediates radiation-induced senescence in cancer cells

**DOI:** 10.1101/2021.02.17.431668

**Authors:** Liat Hammer, Vered Levin-Salomon, Naama Yaeli-Slonim, Moria Weiss, Naama P. Dekel-Bird, Tsviya Olender, Ziv Porat, Sabina Winograd-Katz, Alon Savidor, Yishai Levin, Shani Bialik, Benjamin Geiger, Adi Kimchi

## Abstract

Radiation therapy can induce cellular senescence in cancer cells leading to short-term tumor growth arrest, yet increased long-term recurrence. To better understand the molecular mechanisms involved, we developed a model of radiation-induced senescence in cultured cancer cells, which exhibited a typical senescent phenotype, including upregulation of p53 and its target p21, followed by sustained reduction in cellular proliferation, changes in cell size and cytoskeleton organization, and senescence-associated beta-galactosidase activity. A functional siRNA screen using a cell death-related library identified the mitochondrial Ser protease HtrA2 as necessary for senescence development. Mass spectrometry-based proteomic profiling of the senescent cells indicated downregulation of proteins involved in cell cycle progression and DNA repair, and upregulation of proteins associated with malignancy, while irradiation with HtrA2 inhibition upregulated cell proliferation components. In search of direct HtrA2 substrates following radiation, we determined that HtrA2 cleaves the intermediate filament protein vimentin, affecting its cytoplasmic organization. Ectopic expression of active cytosolic HtrA2 resulted in similar changes to vimentin filament assembly. Thus HtrA2, contributes to several hallmarks of senescence and is involved in the cytoskeletal reorganization that accompanies radiation-induced senescence.

**Summary:** Here the authors identify the Ser protease HtrA2 as a novel mediator of radiation-induced senescence, necessary for sustained proliferation arrest and reorganization of the vimentin filament network.

## Introduction

Radiation therapy is often used to induce cytotoxic responses such as apoptosis in tumors, particularly in patients with non-metastatic lung cancer (Heinzerling et al., 2011), but can also lead to survival responses that avoid cell death, such as autophagy and cellular senescence (Patel et al., 2020). Senescent cells are proliferation arrested, and their cytostatic nature precludes further tumor growth. Yet, it is now recognized that at times, the dormant senescent tumor cells can eventually evade growth arrest and regain their proliferative capacity, eventually leading to tumor recurrence (Saleh et al., 2019). In fact, the presence of senescence markers in patients receiving chemo-radiation therapy has been associated with poor prognosis (Rau et al., 2003). On the other hand, combining conventional treatment with anti-senolytic drugs, such as those that inhibit Bcl2 family members, can block tumor progression (Saleh et al., 2020; Shahbandi et al., 2020).

Cellular senescence can be a natural consequence of aging and telomere shortening, (i.e. replicative senescence), or of cell stress induced by certain oncogenes, chemotherapeutic drugs or radiation (Hinds and Pietruska, 2017). Proliferation-arrested, senescent cells remain metabolically active and have distinct morphologic and biochemical profiles. Their characteristic features include enlarged area, flattened morphology, a prominent nucleus, increased cytoplasmic granularity, accumulation of heterochromatin foci, and enlargement of the lysosomal fraction including enhanced β-galactosidase activity (known as senescence-associated β-Gal, SA-β-Gal) (Ewald et al., 2010). At the molecular level, senescence results from elevated expression of cyclin-dependent kinase inhibitors such as p21 (*CDKN1A*) and p16 (*CDKN2A*) (Ewald et al., 2010). This leads to suppression of E2F transcription factors that drive expression of genes involved in DNA replication and the G1/S and G2/M transitions (Bringold and Serrano, 2000). DNA damage-induced p53 activation and subsequent increased transcriptional activity is a main regulator of p21 expression. As *CDKN2A* is commonly mutated or deleted in cancer, the CDKN2A-RB1 axis is less relevant to the induction of senescence in tumor cells.

In order to better understand the contribution of therapy-induced senescence to the suppression of tumor growth on one hand, and to progression and tumor relapse on the other, a more thorough elucidation of its molecular basis is mandated. Although the DNA damage response, p53 up-regulation and cell cycle arrest pathways are well established, very little is known about the molecular mechanisms governing other aspects of senescence, including the characteristic morphologic changes. Here, we addressed this issue in a model of radiation-induced senescence of the non-small cell lung cancer cell line NCI-H460 by applying an siRNA functional screen and characterizing the proteomic profile by mass spectrometry. We used an siRNA sub-library targeting genes from the cell death and autophagy pathways, since senescence and cell death are alternate fates triggered by the same stressors, with molecular cross-talk among the pathways. We thus identified HtrA2, a mitochondrial Ser protease that is released to the cytosol upon cell stress, as an effector driving the re-organization of the vimentin intermediate filament (IF) network, mediating cytoskeletal alterations that occur during cellular senescence.

## Results

### Characterization of radiation-induced senescence in cancer cells

As a model for therapy-induced senescence, NCI-H460 non-small cell lung cancer cells, which express *TP53* but not *CDKN2A*, were irradiated by clinically relevant doses of X-rays (10 Gy). 24 to 48h post-irradiation, morphologic hallmarks of senescence were observed by phase contrast microscopy: most of the irradiated cells were enlarged and flattened, and in contrast to untreated cells, the cells failed to reach confluency (Fig. S1A). This senescent morphology was observed for at least 14d. The number of viable, metabolically active cells, as measured by PrestoBlue assay, started to decline at 24h compared to untreated cells, and continued to decrease with time (Fig. 1A). Quantitation of live cells indicated that the total number of cells dropped dramatically compared to control untreated cells, which continued to proliferate, but did not significantly change over the 72h time-course (Fig. 1B, Fig. S1B). Thus, the large reduction in PrestoBlue values over time most likely does not reflect loss of cells, but rather a decline in their reductive capacity. Staining with calcein/propidium iodide (PI) to measure live/dead cells at 48h indicated a low percent dead cells, which could not account for the large decreases in PrestoBlue values and cell number compared to control (Fig. S1A). Furthermore, no caspase-3 cleavage was evident, and no subG1 DNA was observed, suggesting that apoptosis was not activated in these cells (Fig. S1C,D). Thus, the reduction in cell number compared to control cells most likely reflects a block in proliferation. Cell cycle arrest at the G1 and G2 phases was confirmed by FACS analysis for DNA content: at 24h post-irradiation, cells accumulated at the G1 and G2/M phases at the expense of S phase and did not incorporate bromodeoxyuridine (BrdU), indicative of a block in cell cycle progression (Fig. S1D). Staining for SA-β-Gal and quantification by ImageStreamX flow cytometry indicated an increase in the portion of irradiated cells that were positively stained at 48h and 72h post-irradiation, compared to the control cells (Fig. 1C). Clear differences were also observed when comparing the mean cell areas (Fig. 1D), reflecting the increased cell size following irradiation (Fig. S1E). At the molecular level, protein levels of both p53 and p21 increased post-irradiation (Fig. 1E). Altogether, the molecular data, combined with the cell proliferation analysis and observed morphologic changes, indicate that radiation induces sustained senescence in these lung cancer cells. Radiation-induced senescence was also observed in HCT116 colon cancer cells, which displayed reduced proliferation, increased cell size, SA-β-Gal accumulation, and induction of p53 and p21 (Fig. S2).

**Fig. 1.**
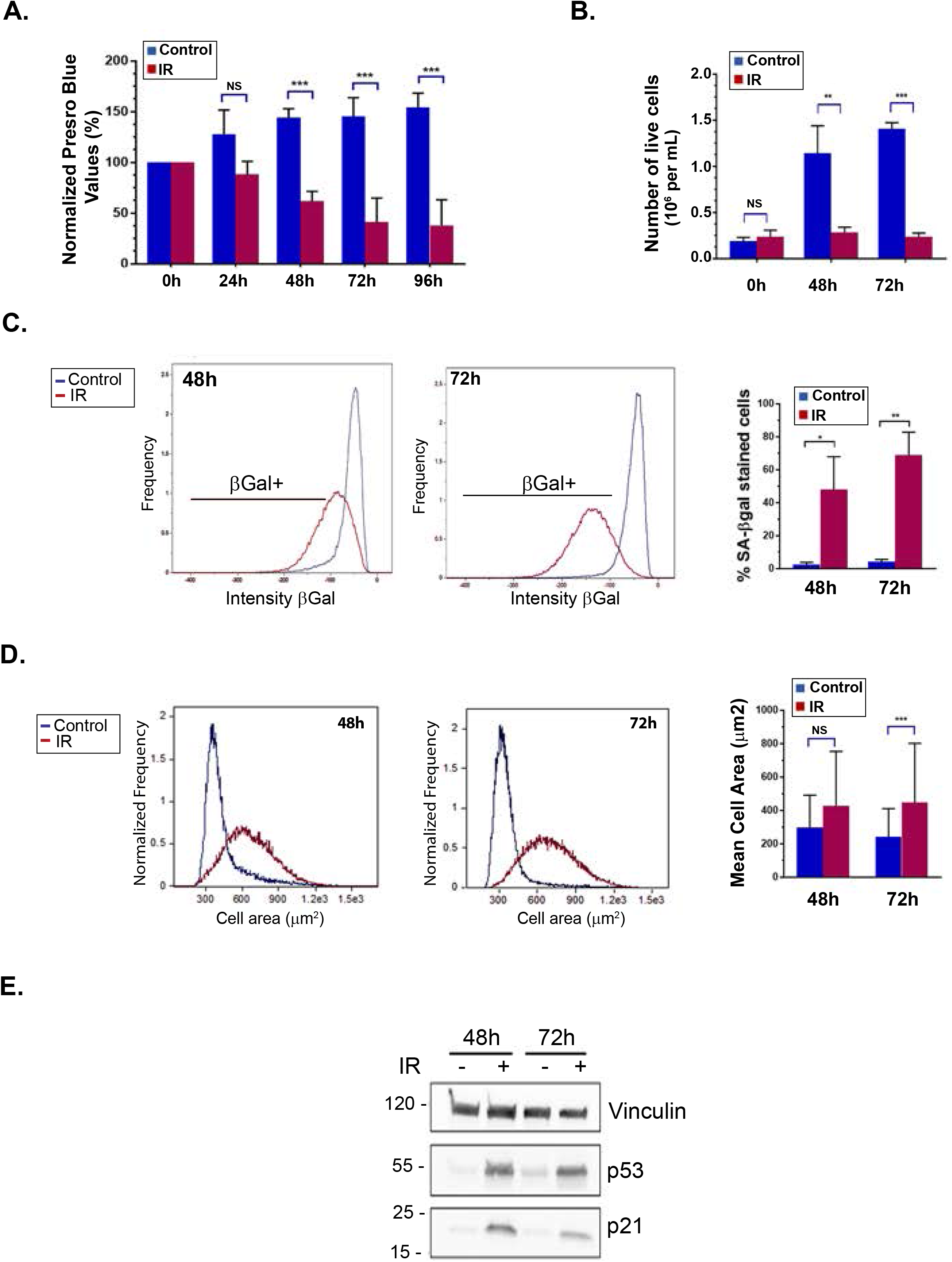
Radiation induces senescence in NCI-H460 lung cancer cells. **A.** Cells were irradiated and cell number/metabolic was measured by PrestoBlue assay at the indicated times post-treatment. Values were normalized to 0h. Shown is the mean±SD of 3 (24h, 48h) or 4 (72h, 96h) biological repeats per time-point. Statistical significance was determined by Student’s two tailed T-test, *** *p* = 0.000497, *p* = 0.000574, *p* = 0.000256, respectively for 48h, 72h, 96h. **B.** Cells were irradiated and live cells counted after the indicated times. Shown is the mean±SD of 4-6 individual counts from a representative experiment. Statistical significance was determined by Student’s two-tailed T-test, ** *p* = 0.00883, *** *p* = 5.63×10^−10^. An additional biological repeat of this experiment can be found in Fig. S1B. **C,D.** 48h or 72h post-irradiation, cells were stained for SA-β-gal and DAPI, followed by ImageStream X analysis. At least 2.9×10^4^ cells were collected from each sample. Representative distributions indicating intensity of SA-β-Gal staining (C) and cell size (D) are shown. Graph on right in C shows the percentage of positively stained cells, as mean±SD of 5 (48h) or 4 (72h) separate experiments. Statistical significance was determined by two tailed Student’s T-test, * *p*=0.00685, ** *p*=0.00246. Graph in D shows the mean cell area±SD of 5 (48h) or 4 (72h) separate experiments. Statistical significance was determined by two tailed Student’s T-test, * *p*=0.0164, ** *p* = 0.00906. **E.** Western blots of lysates from cells 48h or 72h post-irradiation.

### siRNA functional screen targeting genes within the programmed cell death pathway identifies HtrA2 as a positive mediator of radiation-induced senescence

Having established the senescence phenotype that develops in response to radiation of NCI-H460 and HCT116 cells, we next applied a functional siRNA-based screen for discovering new players that regulate/execute the process. Since a cell’s life and death decisions are intricately linked, many points of molecular crosstalk exist within the programmed cell death (PCD) network, and many proteins have dual functions within these various life and death pathways (Bialik et al., 2010). Senescence and cell death are alternative cell fates that are often triggered by the same stressor, and molecular switches can control whether one or the other is activated (Shahbandi et al., 2020; Yosef et al., 2016). Consequently senescence can be considered an arm of the global cell death/viability network. Thus we hypothesized that radiation-induced senescence may be regulated and/or executed by some of the same proteins regulating PCD. In order to identify such dual function players, a small-scale siRNA mediated screen was performed in NCI-H460 cells. The siRNA library, comprising 97 siRNAs targeting apoptosis, programmed necrosis and autophagy genes, and also several known regulators of senescence, was applied to both non-irradiated and irradiated NCI-H460 cells. These were then assayed by CellTiter-Glo (CTG), as a measure of metabolic activity/cell number, to identify PCD genes whose knock-down (KD) affected the senescence response to radiation (Fig. 2A). A reduction in the response following gene KD (i.e., increased CTG values in irradiated cells) indicates that the corresponding gene functions as a positive mediator of cellular senescence, while enhancement of the response (i.e. decreased CTG values) suggests that either the corresponding gene inhibits senescence, or prevents cell death. Fig. 2B shows the fold-change in CTG levels, normalized to non-targeting siRNA control, across the PCD library (see Table S1 for raw and normalized values). The change in CTG was statistically significant for 18 genes (Fig. 2B, red points). The KD of two of these (*BCL2L1* (BCL-XL) and *CRADD*) resulted in an even greater reduction in CTG activity compared to control (Fig. 2C), the former consistent with previous reports on the senolytic effect of BCL-XL KD or inhibition (Saleh et al., 2020; Shahbandi et al., 2020; Yosef et al., 2016; Zhu et al., 2015). The KD of the remainder attenuated the response; those with a > 1.5-fold increase in CTG readout are also listed in Fig. 2C. The top hit that passed the FDR correction threshold was *HtrA2* (Fig. 2C). KD of *HtrA2* resulted in a nearly two-fold increase in CTG values following irradiation, normalized to control KD, i.e. the radiation-induced decrease in cell number/metabolic activity was attenuated by depletion of *HtrA2*. This suggests that HtrA2 (also known as Omi), a mitochondrial Ser protease that is released to the cytosol during cell stress (Vande Walle et al., 2008), may be a positive mediator of radiation-induced senescence in NCI-H460 lung cancer cells.

**Fig. 2.**
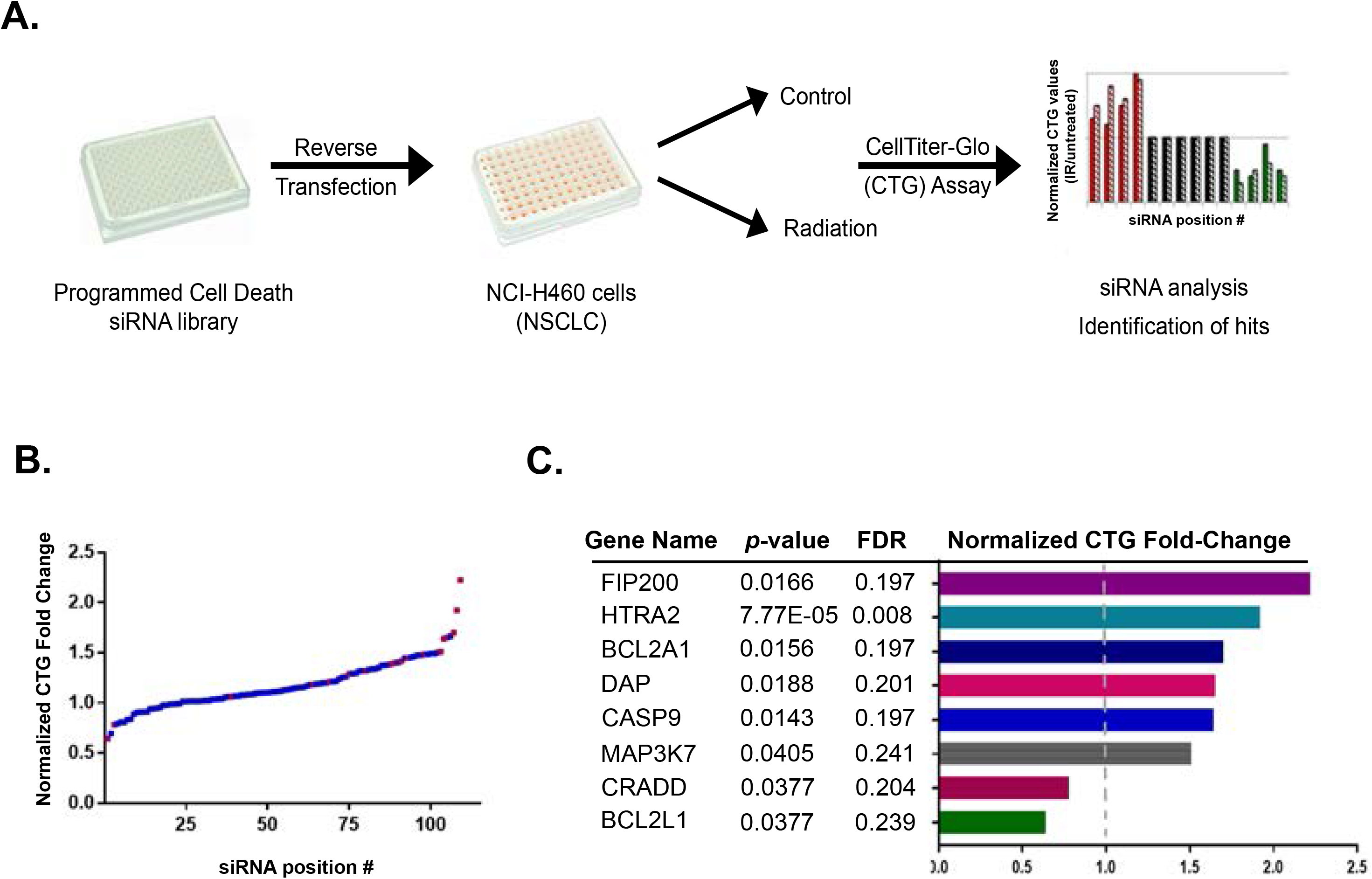
Functional siRNA screen for regulators of radiation-induced senescence. **A.** Schematic of screen using a PCD siRNA library to identify gene KDs that either enhance or attenuate the cellular senescent response to radiation, assessed by CTG to quantitate changes in metabolic activity/cell number. **B.** Distribution of senescence response of siRNA KDs across the library. For each siRNA, the ratio of CTG values for irradiated vs. untreated cells was calculated. Normalized ratios are plotted for each siRNA, with control non-targeting siRNA values set as 1. siRNAs are distributed along the X-axis in increasing order of the normalized ratios; some were assayed twice as internal plate controls and both outcomes are plotted. Red points indicate siRNAs with statistically significant fold change, *p*<0.05. **C.** List of genes showing a significant fold-change in the normalized CTG values. Statistical parameters are indicated, significance was determined by paired T-test. For proteins with increased ratio, only those with a fold-change >1.5 are listed.

Western blot analysis confirmed reduced HtrA2 expression upon introduction of siRNA against *HtrA2* in NCI-H460 cells (Fig. 3A). Consistent with the screen results, microscopy-based examination of irradiated cells indicated that *HtrA2* KD cells were increased in number, reaching near confluency following irradiation, similar to non-irradiated cells (Fig. 3B). The total number of cells (Fig. 3C) and the PrestoBlue values (Fig. 3D) were enhanced by *HtrA2* KD following irradiation, compared to control KD. KD of *HtrA2* also mitigated the radiation-induced increase in SA-β-gal staining compared to irradiated cells transfected with control siRNA (Fig. 3E). Notably, p53 induction was not affected by *HtrA2* depletion (Fig. 3A). In order to evaluate the contribution of HtrA2 to long-term senescence, shRNA against *HtrA2* was used to establish stable KD in NCI-H460 cells (Fig. 3F). Similar to transient KD cells, the stable *HtrA2* KD cell line exhibited reduced SA-β–Gal staining following irradiation, compared to control KD cells, further validating the outcome of *HtrA2* depletion by a different KD strategy targeting different sequences (Fig. S3A,B). Significantly, while irradiated control KD cells maintained reduced PrestoBlue values even at 7d following irradiation, the irradiated *HtrA2* KD cells showed a mitigated senescent response, with increased PrestoBlue values. (Fig. 3F). Finally, to show the relevance of HtrA2 as a regulator of senescence in other cancer cell lines, HCT116 cells were treated with an inhibitor of HtrA2 catalytic activity, Ucf-101, and irradiated. Senescence was measured 48h later by SA-β-gal staining. Significantly, the number of β-gal positive cells decreased in irradiated cells upon HtrA2 inhibition, compared to control treated cells (Fig. 3G). This suggests that HtrA2 is a general positive mediator of radiation-induced senescence in cancer cell lines.

**Fig. 3.**
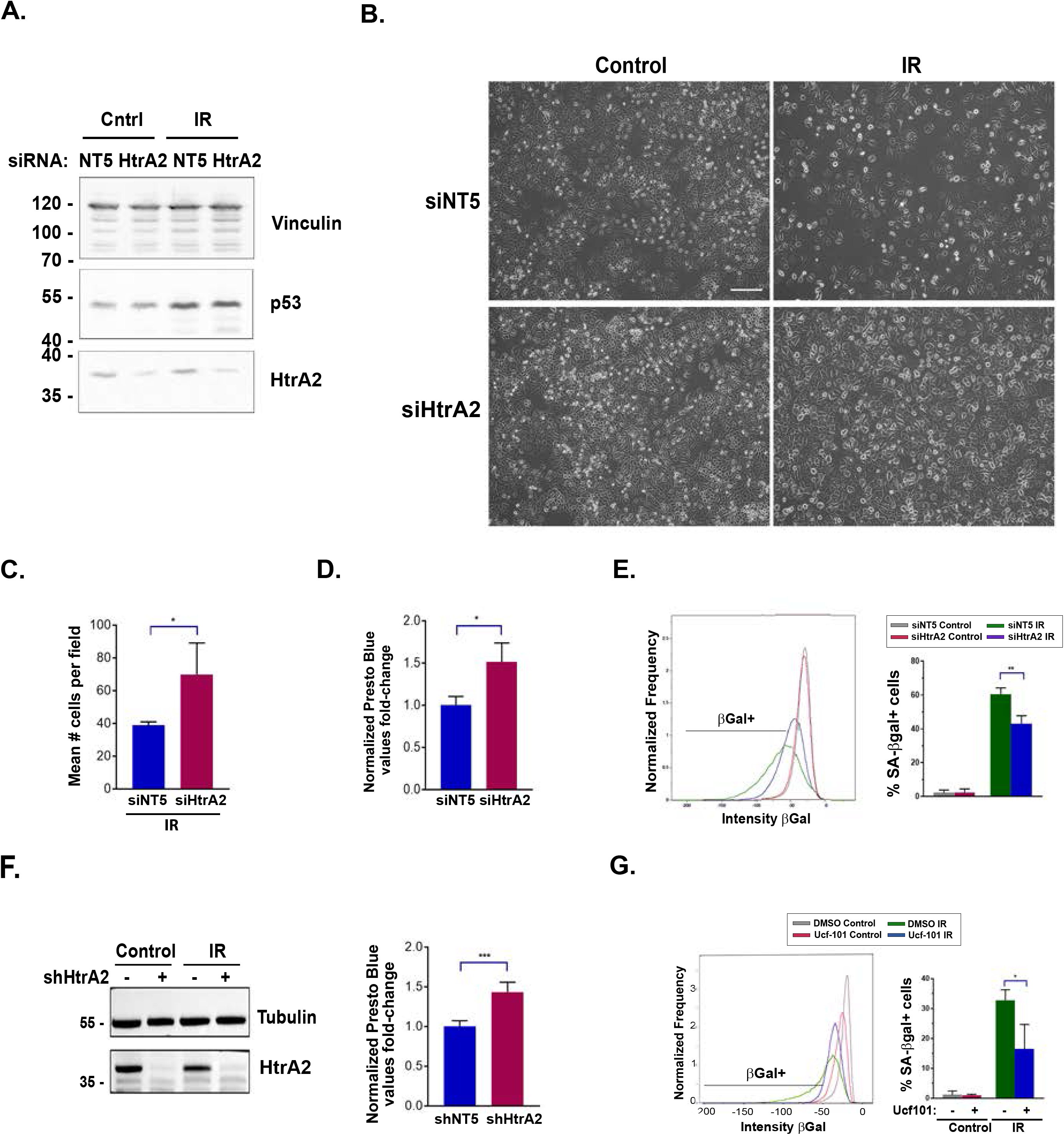
KD or inhibition of HtrA2 blocks radiation-induced senescence in NCI-H460 and HCT116 cancer cells. **A.** NCI-H460 cells expressing siRNA to HtrA2 or control siRNA were irradiated and western blotted 48h later with antibodies to HtrA2 and p53. Vinculin was used as a loading control. **B.** Light microscopic images of control and HtrA2 KD cells 48h post-irradiation. Bar, 200μ. **C.** Cells expressing HtrA2 or control siRNAs were irradiated and the total number of viable cells counted 48h later. Shown is mean±SD of 3 biological repeats, statistical significance determined by two tailed T-test, * *p* = 0.0491. **D.** Fold-change in PrestoBlue values (irradiated/non irradiated) 48h post-treatment upon KD of HtrA2 by siRNA, with siNT values normalized to 1. Shown is mean±SD of 3 biological repeats, statistical significance determined two tailed T-test, * *p* = 0.0235. **E**. siHtrA2 or control NT cells were irradiated and stained for SA-β-gal and DAPI, followed by ImageStream X analysis after 72h. At least 1.3×10^4^ cells were collected from each sample. A representative distribution is shown, and graph represents mean±SD of 3 experiments. Statistical significance was determined by Student’s two tailed T-test, ** *p* = 0.00743. **F.** NCI-H460 cells stably expressing shRNA to HtrA2 or control non-targeting shRNA were irradiated and 7d later, subjected to western blotting to confirm KD and PrestoBlue assay to measure viable cell number/metabolic activity. Graph shows mean fold-change (irradiated values divided by control values)±SD of 4 technical repeats from 1 of 3 representative experiments, normalized to NT control KD, which was set as 1. Statistical significance was determined by Student’s two tailed T-test, *** *p* = 0.000889. **G.** HCT116 cells treated with DMSO or Ucf-101 were irradiated and stained for SA-β-gal and DAPI after 48h. At least 2.6×10^4^ cells were collected from each sample and analyzed by ImageStream X. A representative distribution of SA-β-gal positive cells is depicted, and graph represents mean±SD of 3 experiments. Statistical significance was determined by Student’s two tailed T-test, * *p* = 0.0336.

### Proteomics profiling of irradiated NCI-H460 cells

In order to broaden our understanding of the signaling pathways governing radiation-induced senescence, to characterize alterations in the proteomic profile induced by radiation, and to study how HtrA2 may be connected to these pathways and changes, mass spectrometry was performed on untreated and irradiated NCI-H460 cells, with or without the HtrA2 inhibitor Ucf-101. A total of ~8000 proteins were identified. There was a close correlation within the 4 biological repeats of each experimental setting, with irradiated samples more closely related to each other, and similarly untreated samples, regardless of drug treatment (Fig. S4A). Quantitative statistical analysis of irradiated vs. control DMSO treated samples revealed differential expression of 317 proteins (*p* and *q* values < 0.05, fold change > 2 or <0.5), the abundance of which increased for 170 proteins, and decreased for 147 proteins (Fig. 4A,B and Table S2). As expected, p53 and p53-response genes were among the proteins with increased abundance (24 total, 14.1% of upregulated proteins), including p21, while 107 of the proteins that decreased in abundance (72.8%) were reported targets of p53/p21-mediated DREAM complex transcriptional repression (Engeland, 2018; Fischer, 2017). Thus overall, although not excluding translation regulation and post-translation effects, p53-dependent transcriptional regulation potentially accounts for 41% of the up- and down-regulated genes in the dataset, most likely through direct up-regulation of *CDKN1A*.

**Fig. 4.**
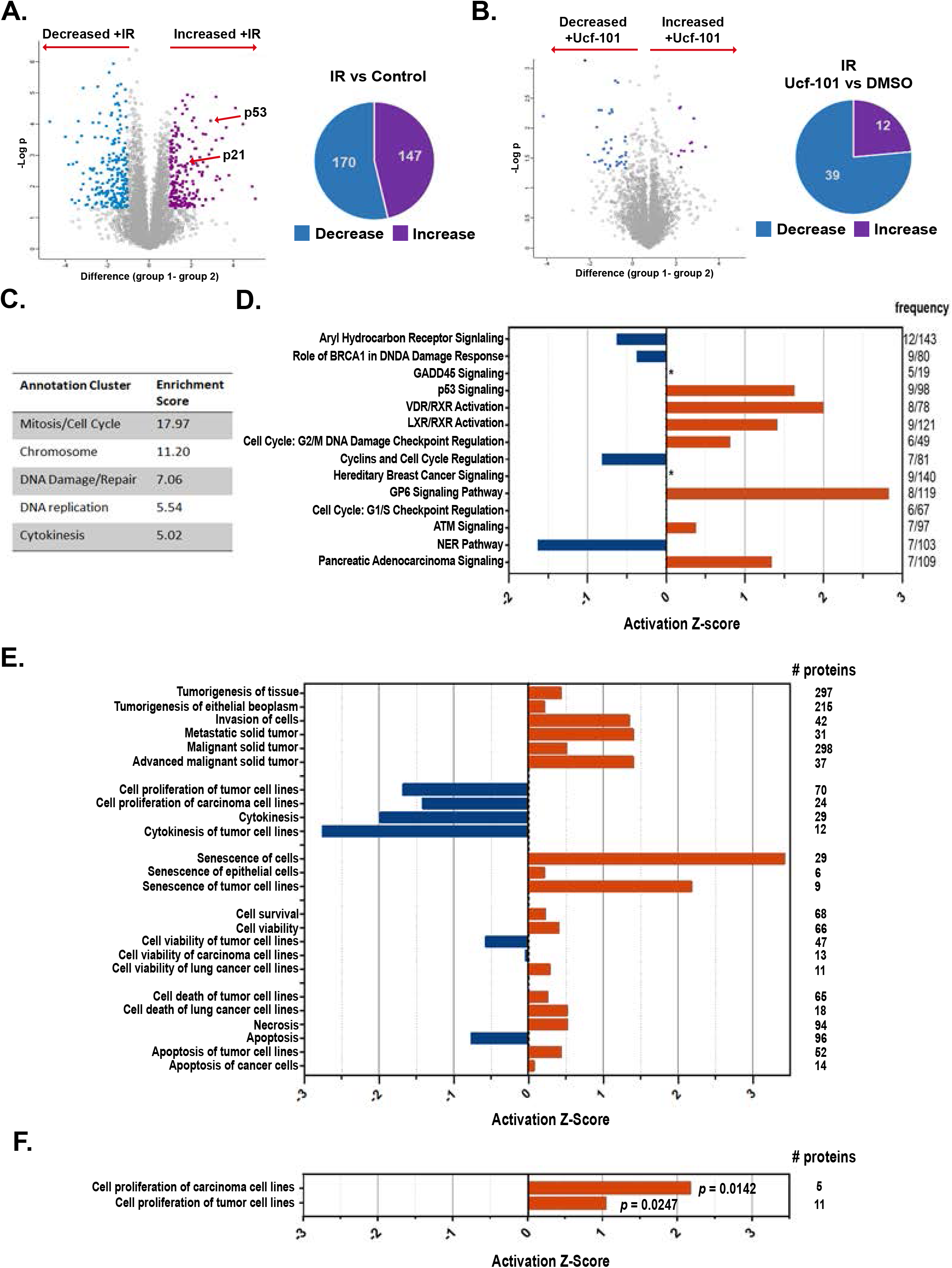
Mass spectrometry analysis of irradiated vs. control non-irradiated cells, and upon HtrA2 inhibition. **A,B.** Volcano plots (left) based on protein LFQ-MS intensity values. Highlighted data points indicate significantly higher (purple) or lower (turquoise) abundance proteins in irradiated vs control cells (A) and in DMSO vs Ucf-101 treated irradiated cells (B), as summarized in pie charts at right. **C**. Gene ontology annotation cluster analysis by DAVID of proteins with altered abundance upon irradiation; top 5 clusters are indicated. **D.** Ingenuity Pathway Analysis showing the top canonical pathways, listed in descending order, of the dataset of proteins with altered abundance following irradiation. Frequency is defined as the number of proteins in dataset/total number of proteins in the given pathway. A positive Z-score indicates likely pathway activation. * Z-score undetermined. **E,F.** Ingenuity Diseases and Functions analysis of the dataset of proteins with altered abundance following irradiation (E) and irradiation with Ucf-101 treatment (F). Highly significant (p < 10^−5^) pathways are shown, grouped by common category. Values on the right refer to the number of proteins in dataset within the given pathway. *p*-values are shown in F.

Gene ontology analysis by DAVID software of this set of proteins revealed that, as expected, the top annotation clusters were related to cell proliferation, including mitosis/cell cycle/cell division, DNA replication, cytokinesis, and DNA condensation (e.g. centromere associated proteins, Aurora Kinase B, kinesin family members, condensin complex subunits, DNA polymerase α, DNA topoisomerase 3α), all with reduced abundance, as well as those involved in cell cycle arrest (e.g. p21 (CDKN1A) and p27 (CDKN1B), both increased in abundance) (Fig. 4C). DNA damage/repair cluster was also highly affected by irradiation; most of the proteins in this category were reduced in abundance, and of those, the majority was identified as p53 repressed gene targets (Engeland, 2018; Fischer, 2017) (Fig. S4B). The dataset was also annotated by Ingenuity Canonical Pathway and Disease and Function analyses, which provide information as to the likely activation state of a given pathway or function (i.e. a positive Z-score implies a positive correlation with the activated state). As expected, G1/S and G2/M checkpoints and senescence functions were predicted to be activated, but in contrast, pathways driving cell cycle progression, cell proliferation and cytokinesis functions were predicted to be repressed (Fig. 4D,E). While p53 and ATM signaling are likely activated, the BRCA1-related DNA-damage response is likely to be inactivated (Fig. 4D), consistent with the DAVID gene ontology analysis. Interestingly, malignancy functions featured prominently among the biological processes with high positive Z-scores, including pathways such as tumorigenesis, invasion, and metastasis (Fig. 4E). The Ingenuity classifications related to cell death and survival were less informative; cell viability and cell survival functions had both low positive and low negative Z-scores, making these results more difficult to interpret. Of note, the proteins within these categories did not include any of the central regulators or executioners of PCD. Overall, the proteomics analysis of irradiated NCI-H460 cells was consistent with the induction of senescence, but not cell death, with a diminished DNA damage response and surprisingly, up-regulation of proteins associated with malignancy.

Comparison of the sets of proteins with altered expression in Ucf-101 treated vs. DMSO treated irradiated cells yielded a smaller group of 51 proteins, 12 of which increased in abundance and 39 decreased (Fig. 4B and Table S3). Ingenuity Disease and Function analysis indicated activation of proliferation functions (Fig. 4F), which contrasted with the predicted inactivation of these pathways by irradiation alone (Fig. 4E). Thus, the effect of HtrA2 inhibition on the proteomic signature of the irradiated cells is consistent with the observed mitigation of the sustained proliferative block in response to radiation shown in Fig. 3. Notably, p53, and p53-target genes that were up-regulated by irradiation showed no significant change upon HtrA2 inhibition. Likewise, the majority of the 107 p53 repression targets that were reduced following irradiation were not affected by the addition of the HtrA2 inhibitor, with the exception of 2 targets whose abundance increased (RAD54B, DHCR7) and 3 others that showed an even greater reduction in the presence of Ucf-101 (CENPH, ELF1, HIRIP3). Thus, consistent with the western blotting showing persistent induction of p53 and p21 following irradiation even upon HtrA2 KD or inhibition (Fig. 3A, Fig. S3C), the MS data suggest a role for HtrA2 downstream to p53 function.

### HtrA2 cleaves vimentin and affects vimentin assembly and polymerization following irradiation

To investigate the p53-independent function of HtrA2 in senescence, we focused on identifying HtrA2 cytosolic substrates, as previous work demonstrated that it is released from the mitochondria into the cytosol following cell stress (Martins, 2002). HtrA2 was shown to cleave the IF vimentin *in vitro*, resulting in the removal of part of its N-terminal head domain, yielding a truncated protein of 49 kDa (Vande Walle et al., 2007). Therefore, vimentin expression was assessed here by western blot to determine if it is cleaved in senescent cells. As shown in Fig. 5A,B, a faster migrating form of vimentin was observed following irradiation in NCI-H460 cells, consistent with the proteolytically cleaved form of vimentin (Lucotte et al., 2015). The levels of this smaller form of vimentin was reduced in HtrA2 KD cells (Fig. 5A) and upon treatment of cells with Ucf-101 to inhibit HtrA2 (Fig. 5B). Thus, cleavage of vimentin during radiation-induced senescence requires HtrA2 proteolytic activity.

**Fig. 5.**
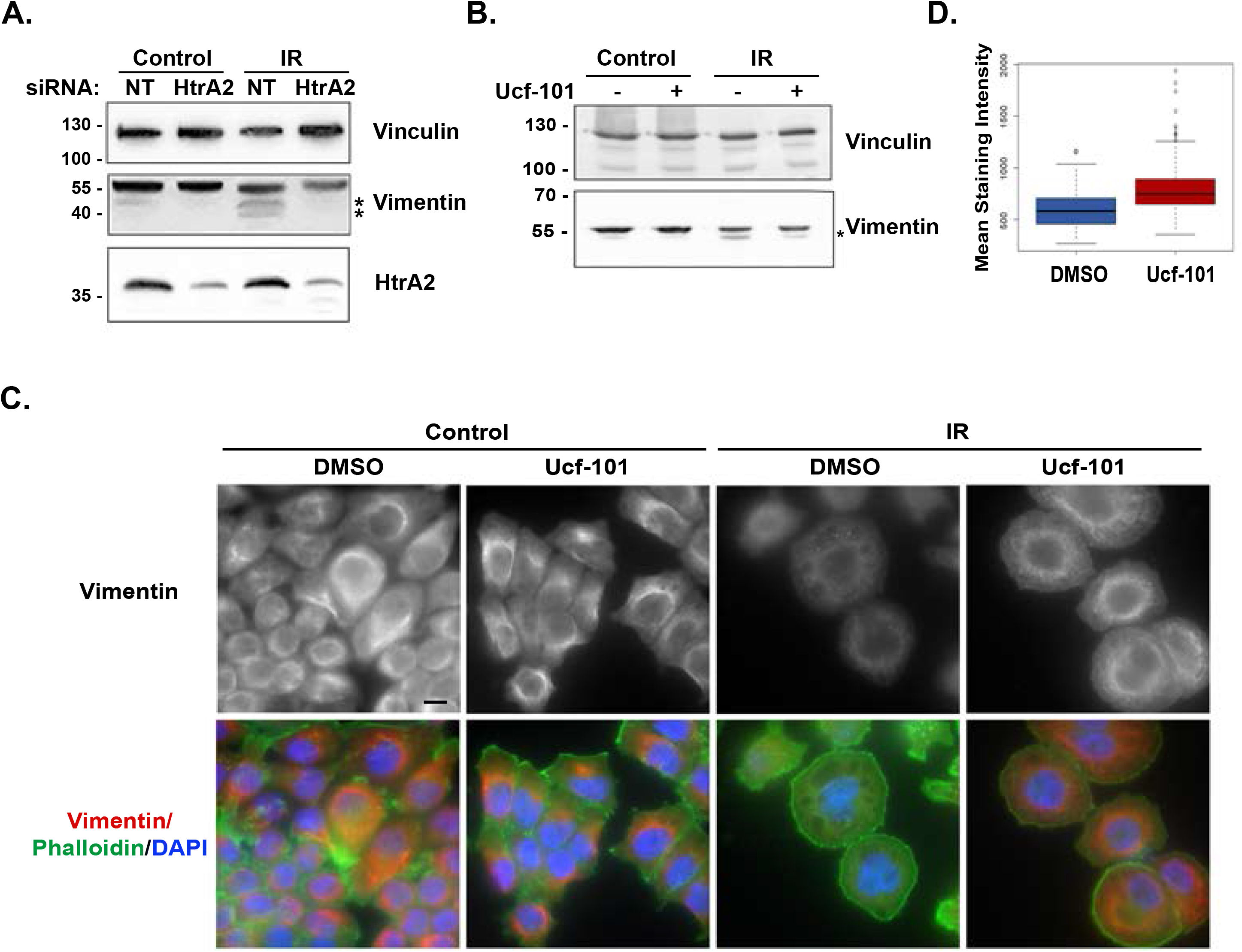
Radiation-induced cleavage of vimentin and decreases in vimentin staining intensity require HtrA2 proteolytic activity. **A., B.** NCI-H460 cells expressing HtrA2 or control siRNA (A) or treated with either Ucf-101 or DMSO as control (B) were irradiated and western blotted 48h later with antibodies to HtrA2 or vimentin. Asterisks indicate vimentin cleavage products. Vinculin was used as a loading control **C.** Cells treated with either DMSO or Ucf-101 were irradiated and 48h later immunostained for vimentin. Phalloidin (green) and DAPI (blue) staining was used to detect actin and nuclei, respectively, and are shown in merged images at bottom. Note that all images are the same magnification, bar 10μ. **D.** Box plot showing the range of vimentin staining intensity measured according to the average pixel intensities inside the contours of each cell. Black lines represent median values, colored regions represent the 25th-75th percentiles, and dots represent potential outliers. Statistical significance was determined by Two-sample Kolmogorov-Smirnov test for two biological repetitions, *p* < 2.2e^−16^.

Previous reports documented that the N-terminal head domain of vimentin is essential for vimentin filament assembly (Beuttenmüller et al., 1994; Dave and Bayless, 2014). To determine if vimentin assembly or localization is affected by senescence in an HtrA2 dependent manner, immunostaining for endogenous vimentin was performed in irradiated NCI-H460 cells treated with Ucf-101, which more uniformly achieves HtrA2 inhibition compared to transient siRNA transfection (Fig. 5C). The intensity of vimentin staining was significantly reduced following irradiation. Most importantly, inhibition of HtrA2 activity by Ucf-101 blocked this decline, without affecting the overall shape of the senescent cells (Fig. 5C,D). Notably, no obvious changes were observed in the actin cytoskeletal network upon addition of Ucf-101 within the 48h time frame post-irradiation (Fig. 5C).

A more comprehensive analysis of vimentin organization in the irradiated cells revealed prominent, variable changes in vimentin IF assembly and distribution, which were mitigated by Ucf-101. Most of the irradiated cells (69%) exhibited diffuse distribution of vimentin throughout the cytoplasm, without visible filaments (Fig. 6A), whereas a minority of cells (11%) displayed discrete, well organized filaments in either the perinuclear region or uniformly distributed throughout the cytoplasm. A fraction of cells (20%) displayed a combination of diffuse and discrete filamentous organization (Fig. 6A), probably representing a transient intermediate stage in the senescence-related IF disassembly. These changes in vimentin organization were HtrA2 dependent, as the addition of Ucf-101 to the irradiated cells significantly reduced the fraction of cells displaying the diffuse pattern (17%), and the fraction of cells exhibiting well-organized discrete filaments reached 66% (Fig. 6B).

**Fig. 6.**
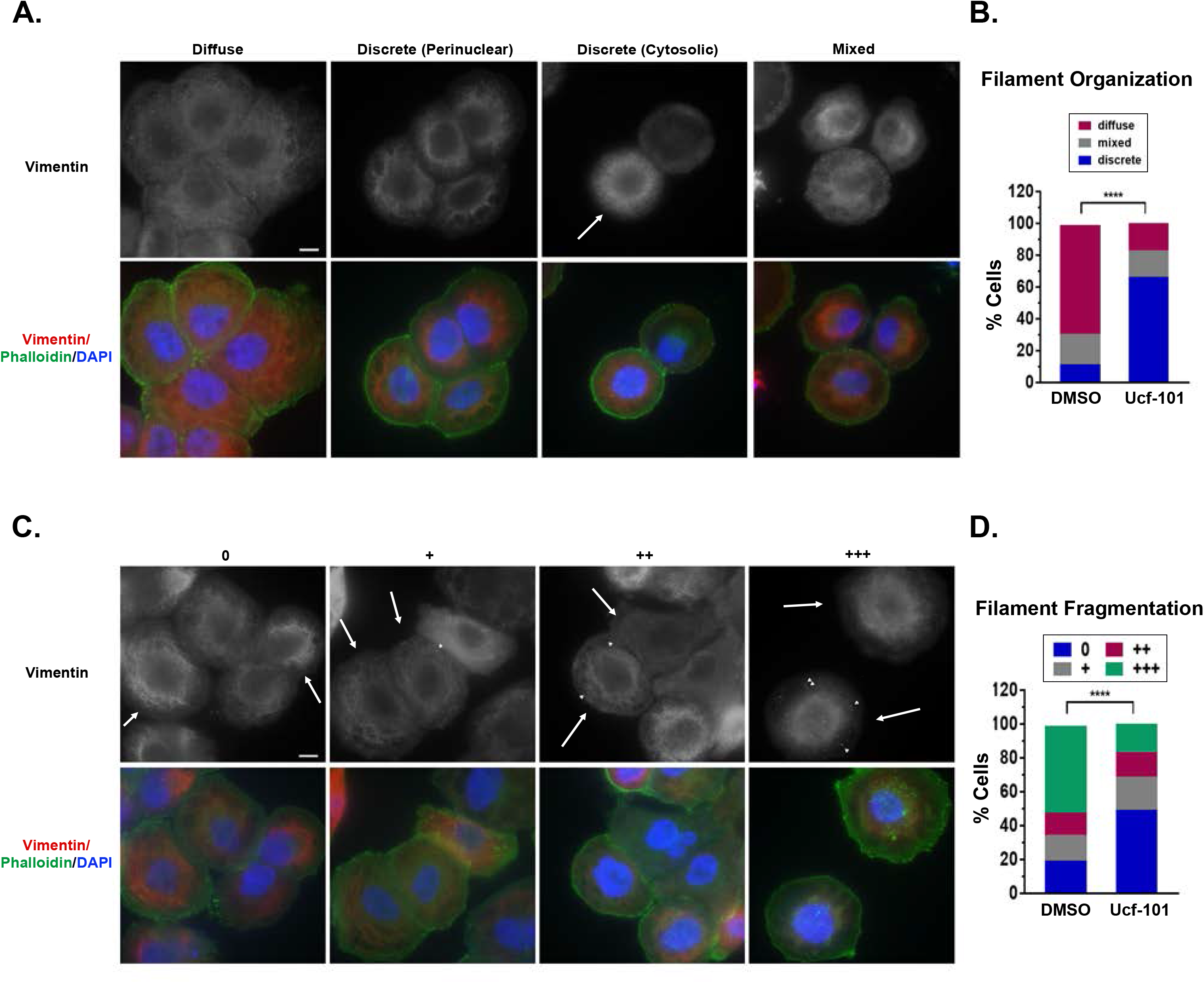
Inhibition of HtrA2 proteolytic activity reduces vimentin filament fragmentation and reorganization following radiation. **A.,B.** NCI-H460 cells were irradiated and 48h later stained for vimentin (red), actin (phalloidin, green) and nuclei (DAPI, blue). Representative images of vimentin stained filaments depicting each type of organization are shown in A. Arrows indicate cells displaying the relevant category, arrowheads, vimentin aggregates. Brightness and contrast were adjusted when needed across an image to improve visibility of specific structures. Bar, 10μ. Quantities of each category in DMSO or Ucf-101 treated cells are depicted in B. Approximately 200 cells were evaluated in each of two biological experiments. Statistical significance was determined by Chi-square test, **** *p* < 0.00001. Cytosolic and perinuclear localized discrete filaments were considered as one category. **C., D.** Same as A and B, but representing degree of filament fragmentation.

In many irradiated cells that did not exhibit discrete vimentin filaments, globular dot-like aggregates were detected. These aggregates are presumed to be fragmented vimentin filaments, as described previously (Beuttenmüller et al., 1994). Cells were visually scored for these aggregates as ‘0’ (no visible aggregates), +, ++, and +++ (increasing amounts of vimentin aggregates), as shown in Fig. 6C. Quantitative analysis of these categories indicated that 52% of irradiated DMSO treated cells contained large amounts of vimentin aggregates, with no visible aggregates in only 19%. In contrast, only 16% of irradiated Ucf-101 treated cells had large number of aggregates, while 49% did not show any dot-like aggregates (Fig. 6D).

To test whether the above mentioned changes in vimentin filament organization is exclusively driven by HtrA2’s presence in the cytosol, NCI-H460 cells were transfected with a plasmid driving the expression of a mature, active HtrA2 lacking the mitochondrial targeting signal and tagged at the C-terminus with GFP (HtrA2Δ133-GFP) (Fig 7A). Transfection with free GFP was used as a control. To avoid apoptosis due to expression of large quantities of cytosolic HtrA2, the pan-caspase inhibitor Q-VD-OPh was added. As expected, HtrA2Δ133-GFP localized to the cytosol in non-irradiated cells (Fig. 7B). The average vimentin staining intensity was significantly lower in the presence of HtrA2Δ133-GFP, compared to GFP control (Fig 7B,C). Furthermore, the majority of HtrA2Δ133-GFP transfected cells exhibited a diffuse distribution of vimentin filaments (70% vs. 14% of GFP transfected cells), with only 8% exhibiting the organized discrete filaments, compared to 65% of GFP expressing cells (Fig 7B,D). Notably, phalloidin staining of HtrA2Δ133-GFP transfected cells indicated that HtrA2 had no obvious effects on the actin cytoskeletal network (Fig 7B). Thus ectopic expression of cytosolic active HtrA2 results in similar changes to vimentin assembly as induced by radiation.

**Fig. 7.**
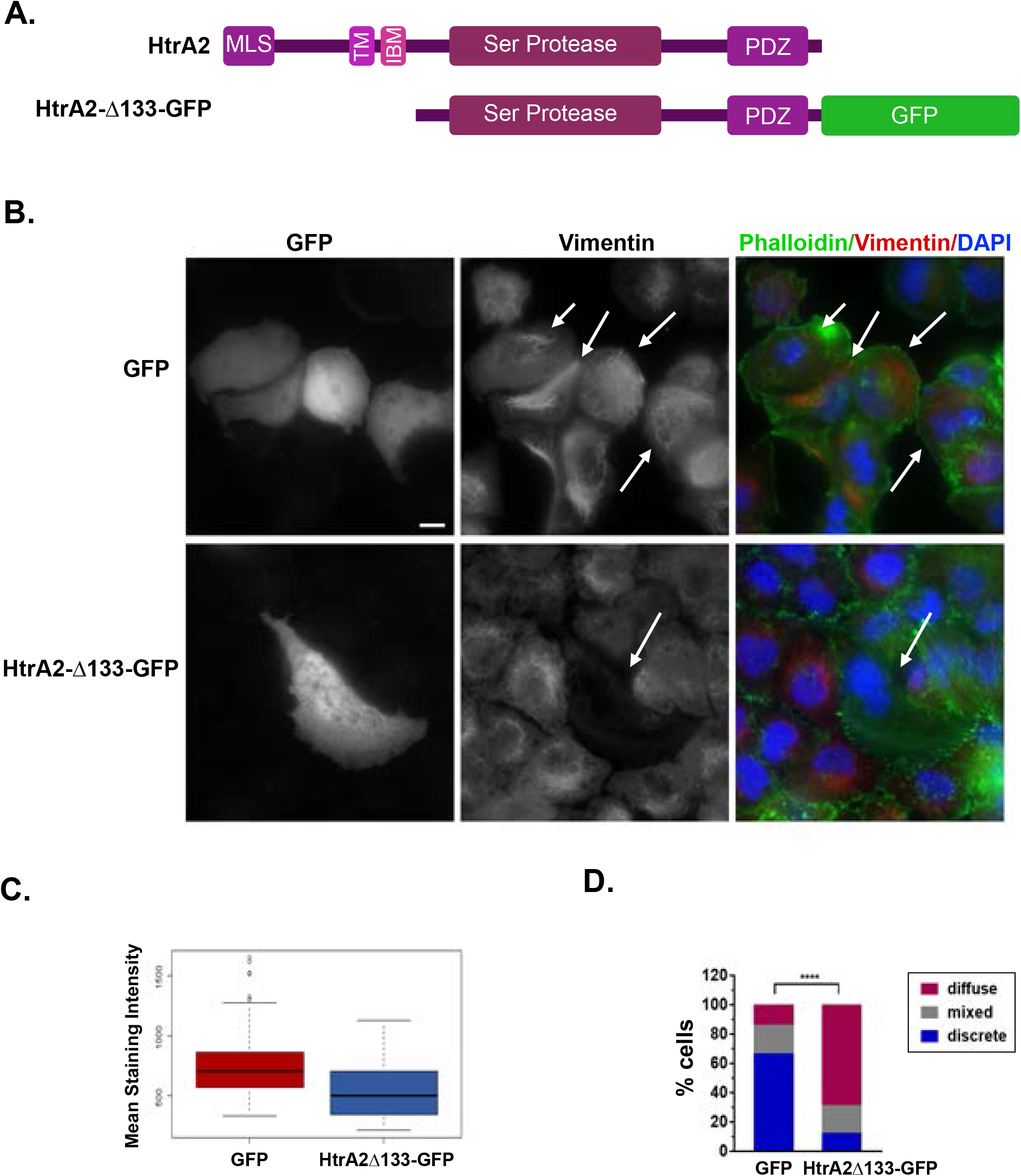
Expression of cytosolic HtrA2 induces changes in vimentin filament organization. **A.** Schematic representation of full length HtrA2 protein, and the cytosolic HtrA2Δ133-GFP construct. MLS, mitochondrial localization signal, TM, transmembrane segment, IBM, IAP binding motif, PDZ, C-terminal PDZ domain. **B.-D.** Cells treated with Q-VD-OPh were transfected with GFP or HtrA2Δ133-GFP. 48h later, cells were stained for vimentin, actin (Phalloidin, green) and nuclei (DAPI, blue). **B.** Representative images of vimentin staining and merged images. Arrows indicate transfected cells. Bar, 10μ. **C.** Box plot showing the range of vimentin staining intensity measured according to the average pixel intensities inside the contours of each cell. Black lines represent median values, colored regions represent the 25th-75th percentiles, and dots represent potential outliers. Statistical significance was determined by Two-sample Kolmogorov-Smirnov test for two biological repetitions, *p*= 1.184e^−06^. **D.** Quantification of vimentin filament organization patterns. Statistical significance was determined by Chi-square test for two biological repetitions, **** *p* < 0.00001.

In conclusion, following irradiation, concomitant with the acquisition of a senescent state, HtrA2 drives a substantial disruption of the vimentin-based IF network, characterized by decreased amount of visible filaments and prominent aggregates, consistent with filament fragmentation, and decreased overall vimentin staining intensity. Inhibition of HtrA2’s catalytic activity attenuated this process, by significantly preventing the changes in the vimentin filament network. Thus HtrA2 is responsible for these specific cytoskeletal changes associated with radiation-induced senescence.

## Discussion

As shown in this study, radiation induces cellular senescence in NCI-H460 lung cancer cells and HCT116 colon cancer cells, as defined by long-term proliferation arrest, changes in cell morphology and size, reduced metabolic activity and enhanced β-gal staining. A functional siRNA screen of PCD proteins aimed to uncover novel regulators of radiation-induced senescence identified HtrA2 as a positive mediator. The contribution of HtrA2 to several aspects of the senescence phenotype, including reduced proliferation and SA-βGal staining, was confirmed in different cancer cell lines, using several KDs approaches, including an siRNA pool and shRNA targeting different sequences, and the specific HtrA2 inhibitor, Ucf-101. HtrA2 is a Ser protease and chaperone that localizes to the mitochondrial inter-membrane space, from where it maintains mitochondrial homeostasis (Vande Walle et al., 2008). In this capacity, its inactivation is associated with neurodegenerative disease and aging (Martins et al., 2004). During cell stress, however, HtrA2 is released to the cytosol as a result of changes in mitochondria (Vande Walle et al., 2008). Within the cytosol, HtrA2 has been shown to induce apoptosis through multiple pathways, including binding to and cleaving IAPs (Martins, 2002; Yang et al., 2003). In the current study, we show for the first time that in addition to its known role in apoptosis, HtrA2 is involved in the regulation of senescence. As apoptosis is not induced in these cells following radiation; either additional factors counter HtrA2’s ability to cleave its apoptotic substrates, or the amounts of HtrA2 released to the cytosol are limiting. In either case, HtrA2 is likely to have alternative targets/substrates that require its protease activity for generating part of the senescence phenotype.

HtrA2 suppression had profound effects on the altered IF organization of the senescent cells, providing for the first time, a mechanistic explanation for cytoskeletal changes associated with senescence. HtrA2 has been shown in the past to cleave several cytoskeletal proteins *in vitro,* including actin and vimentin (Vande Walle et al., 2007), and to modulate G/F-actin dynamics in Ras-transformed senescent fibroblasts by directly cleaving β-actin (Yamauchi et al., 2014). HtrA2 has also been shown to cleave the type III IF vimentin in stressed neuronal cells (Lucotte et al., 2015). While no major changes in the actin filament network were observed following HtrA2 inhibition or overexpression in the current study, HtrA2 proteolytic activity was necessary for vimentin cleavage following radiation in NCI-H460 cells. This cleavage was previously shown to remove the first 40aa of vimentin, resulting in removal of part of the 95aa N-terminal head domain (Vande Walle et al., 2007). Vimentin’s central rod region mediates dimerization and alignment of the dimer into an antiparallel, staggered tetramer, and the exposed head domain interacts with various cellular components, such as lipids, plasma membrane, DNA and RNA. It is also the site of various post-translational modifications, including several phosphorylation events, which affect overall filament assembly. Moreover, the most N-terminal half of the head domain is directly necessary for filament assembly of the tetramers (Herrmann and Aebi, 2004). In fact, we show here that filament assembly is disrupted following irradiation, with increased aggregation and diffuse localization. These aggregates most likely correspond to fragmented vimentin filaments, resembling those described in previous work, in which truncated vimentin deleted of the head domain was introduced into vimentin null cells and led to the formation of vimentin aggregates (Beuttenmüller et al., 1994). These changes in vimentin were blocked by inhibition of HtrA2 catalytic activity, and conversely, forced expression of a proteolytically mature form of HtrA2 in the cytosol was sufficient to induce these changes in vimentin filament distribution and assembly. Although only a small portion of vimentin is cleaved, the functional effect on filament assembly is pronounced. This may suggest that the truncated form acts as a dominant negative to affect assembly of the full length protein. While a previous report showed that expression of high levels of human vimentin lacking the entire head in mouse fibroblasts did not affect endogenous vimentin organization (Andreoli and Trevor, 1994), other reports are more consistent with the dominant negative hypothesis. For example, some vimentin head mutants and truncations failed to form filaments on their own and disrupted wild type endogenous filaments, resulting in the accumulation of aggregates (Beuttenmüller et al., 1994). Interestingly, a mutant variant of vimentin, found in a patient with a premature aging syndrome, promoted the generation of an N-terminal cleaved form, and faster protein turn-over (Cogne et al., 2020). When expressed in vimentin null cells, the mutant failed to from filaments and instead accumulated in cytosolic aggregates, alone or when co-expressed with wild type vimentin. In addition, the cleaved N-terminus fragment, although not observed on western blots due to its small size, may remain stable and interact with the acidic central rod domain of intact vimentin. In fact, the head domain was shown to prevent assembly of vimentin into filaments and unravel already formed filaments *in vitro* (Traub et al., 1992). Altogether, the data suggest that the direct cleavage of vimentin by HtrA2 within the head domain is the main cause for the loss of vimentin filament organization, although indirect effects of HtrA2 that may influence vimentin assembly by other mechanisms are not excluded. In any case, HtrA2’s effects on vimentin assembly underlies the major cytoskeletal change that characterizes senescence in our system.

Although absent in normal epithelial cells, vimentin expression is up-regulated during oncogenic transformation (Satelli and Li, 2011), and is considered a hallmark of the Epithelial-Mesenchymal Transition (EMT) in motile epithelial cells during embryonic development and in metastatic cancer cells (Lowery et al., 2015). Changes in vimentin have also been associated with senescence, although the underlying mechanisms were not explored. For example, doxorubicin-induced senescence in A549 lung adenocarcinoma cells also led to changes in vimentin filament organization (Litwiniec et al., 2010), and vimentin filament organization differed between aging senescent and young, proliferating fibroblasts (Nishio et al., 2001). Notably, vimentin intermediate filaments are involved not only in defining cell structure and shape, but also in cellular motility, organelle positioning, and signal transduction (Lowery et al., 2015). Moreover, it has been shown to be necessary for proliferation and tumor growth (Virtakoivu et al., 2015). Thus, the HtrA2/vimentin axis may control several distinct events during cellular senescence, although additional as yet unidentified HtrA2 substrates may also link it to other molecular pathways mediating the various senescent hallmarks.

To explore such pathways, we conducted a comprehensive mass spectrometric analysis of the proteome content of irradiated cancer cells undergoing senescence in order to derive a specific proteomic signature that defines these senescent cells and to determine how inhibition of HtrA2 affects this signature. To the best of our knowledge, this is the first time such a characterization has been reported in the published literature. It would be interesting to compare this proteomic profile to other cell types, both of normal and cancer origin, and under additional scenarios of stress-induced or replicative senescence, to identify underlying commonalities that define senescence. This may facilitate further mechanistic understanding of the cellular senescence signaling pathway, provide additional molecular markers for monitoring of the process, and potential therapeutic targets to prevent senescence. Interestingly, functions connected to hallmarks of malignancy, including metastasis, tumor invasion and tumorigenesis, were predicted to be activated in the irradiated senescent cells by Ingenuity Diseases and Functions analysis. This finding is counter-intuitive, since it contrasts with the reduced proliferation that was both predicted by the proteomics analysis and observed in the senescent population. However, it is in line with recent clinical outcomes of radiation therapy that suggest that in the long-term, senescence is not a beneficial consequence of cancer therapy. As senescent cells can overcome the mitotic block and regain their ability to proliferate, their presence can lead to tumor regression and resistance to cancer therapy (Krtolica et al., 2001; Saleh et al., 2019). In addition, with time senescent cells develop the senescence-associated secretory phenotype (SASP), which can lead to the development of a pro-inflammatory, immunosuppressive microenvironment that favors tumorigenesis (Hinds and Pietruska, 2017). The expression changes in the dataset that were associated with malignancy, seen early after irradiation, suggest that the senescent cells may already be primed in their current benign state towards this eventuality, even prior to SASP induction.

A relatively small set of proteins changed in abundance upon HtrA2 inhibition in irradiated cells. Significantly, these included proteins linked to proliferation, which was predicted to be activated by HtrA2 inhibition. This is consistent with the enhanced cell proliferation observed upon HtrA2 depletion and/or inhibition. Interestingly, the senescence-associated proliferation block was mitigated despite the apparent continued functionality of the p53/p21 pathway, which is the main mechanism for cell cycle arrest in these *CDKN2A* mutated cells. Both p53 and p21 were upregulated even upon HtrA2 KD and inhibition, and the MS analysis indicated that HtrA2 inhibition did not change the abundance of known p53 target genes nor targets of p53/p21 DREAM-mediated repression. This suggests that HtrA2’s contribution to senescence, including the sustained proliferation arrest, involves a mechanism independent of or downstream to p53 and p21, elucidation of which requires additional investigation.

To conclude, we have identified a novel role for the mitochondrial Ser protease HtrA2, previously implicated in apoptosis, in regulating some of the cytoskeletal changes occurring during radiation-induced senescence. HtrA2 thus joins the short list of proteins, including p53 and BclXL (BCL2L1), which modulate apoptotic cell death and cellular senescence (Borras et al., 2020; Vicencio et al., 2008). *BCL2L1* also emerged in our screen as a gene whose KD enhanced the radiation-induced decrease in cell viability. This molecular crosstalk emphasizes the intricacy by which cellular life and death decisions are linked and opens new avenues for discovery of the molecular regulation of cellular senescence.

## Materials and Methods

### Cell culture and reagents

All reagents were purchased from Sigma-Aldrich unless otherwise indicated. NCI-H460 and HCT116 cells were purchased from ATCC and routinely tested for mycoplasma. Cells were cultured in RMPI-1640 medium (Biological Industries, Beit Haemek, Israel), supplemented with 10% FBS (GibcoBRL), 4 mM glutamine (GibcoBRL), and combined antibiotics (100 μg/ml penicillin, 0.1 mg/ml streptomycin). NCI-H460 shRNA transfected cells were grown in the presence of puromycin (3μg/ml). Cells were treated with the following reagents: DMSO, TRAIL (100ng/ml, PeproTech), Ucf-101 (20 μmol/L, Merck), Q-VD-OPh (50μM, Merck). For radiation experiments, NCI-H460 (8×10^5^ cells) or HCT116 (4 ×10^6^ cells) cells were irradiated 24h after plating by X-rays at 10Gy or 8Gy, respectively, in a single fraction, using XRAD 320 (Precision X-Ray). Control cells were subjected to mock irradiation. When indicated, Ucf-101 or DMSO were added 3h prior to irradiation.

### Cell viability assays

The number of live cells was determined by Countess cell counter (ThermoFisher) or manually by averaging the number of cells in 4 1.3×1.75mm^2^ fields for each 10cm plate under light microscopy. For PrestoBlue assays, PrestoBlue cell viability reagent (Invitrogen, USA) was added to culture medium according to the manufacturer’s protocol. Fluorescence was measured with a microplate fluorescence reader (TECAN, Tecan Trading AG, Switzerland). For CTG assays, luminescence based CellTiter-Glo reagent (Promega, Madison, WI, USA) was added to culture medium according to the manufacturer’s protocol. Luminescence was read in a microplate luminometer (TECAN). For calcein/PI assays, 1μM Calcein AM (Life Technologies) and 1.5μM PI were added to culture medium 24h post-irradiation and live and dead cells were manually counted under light microscopy (Olympus IX73).

### DNA constructs and transfection procedures

GFP was expressed from pEGFP plasmid. pcDNA3-delta133Omi-EGFP was a gift from L. Miguel Martins (Addgene plasmid #14124; http://n2t.net/addgene:14124; RRID:Addgene_14124) (Martins et al., 2002). For transient transfection of DNA, TransIT-X2 reagent (Mirus) was used according to the manufacturer’s instructions. For transient siRNA transfections, 25nM siGENOME siRNA pool to HtrA2 (cat# M-006014-04; see Table S4 for targeted sequences of duplexes in pools) or siCONTROL non-targeting siRNA #5 (NT5, cat# D-001210-05-05) was mixed with Dharmafect1 (Dharmacon), and added to 2×10^6^ cells 24h after plating. For stable shRNA transfections, NCI-460 cells were infected with lentiviral particles containing pGIPZ vector carrying shRNA targeting HtrA2 (Dharmacon clone ID V3LHS_315866, mature antisense sequence: ATAAGGTCAGTGTTTCTCG), GAPDH shRNA as positive control or non-silencing shRNA as negative control, according to the manufacturer’s protocol (Dharmacon, cat# VGH5526). After continuous selection with puromycin, the clone exhibiting the most prominent decreased expression level of HtrA2 was chosen for further analysis.

### Immunostaining and Microscopy

24h post-plating in 35mm glass-bottom microwell plates, NCI-H460 cells were irradiated. After 48h, cells were fixed with 4% paraformaldehyde and 0.1% glutaraldehyde, washed and permeabilized by 0.2% Triton-X100 (Merck), followed by overnight incubation with chicken anti-vimentin polyclonal antibody (Abcam, cat# 24525) in blocking solution of 8% BSA, 0.1% Triton-X100 and then Alexa Fluor 647 conjugated secondary anti-chicken IgG antibody (Invitrogen). Cells were then stained with phalloidin-TRITC and DAPI. Images were acquired with an automated inverted microscope (DeltaVision Elite system IX71 with Resolve3D embedded imaging software; Applied Precision/ GE Healthcare, Issaquah, WA) using a 60x/1.42 oil objective (Olympus, Tokyo, Japan). Further image processing was performed using ImageJ (NIH Imaging Software) (Schneider et al., 2012).

For brightfield light microscopy imaging, cells were viewed by UPLFLN PH 10x/0.30 and 20x/0.50 objectives on an Olympus IX73 fluorescent microscope equipped with a DP73 camera or an Olympus IX71 fluorescent microscope equipped with a DP70 camera. Images were captured with Olympus CellSens software.

### SA-β-gal staining and Imagestream X analysis

SA-β-gal staining protocol and ImageStream X analysis were previously described (Biran et al., 2019). Briefly, irradiated cells were fixed in 4% PFA and stained for SA-β-gal and DAPI. Samples were analyzed by Amnis ImageStream X Mark II (Luminex, Austin, Texas), using dedicated image analysis software (IDEAS 6.2). Cells were gated for single cells according to their area (in μm^2^) and aspect ratio (the Minor Axis divided by the Major Axis of the best-fit ellipse) of the brightfield image. To eliminate out of focus cells, cells were gated using the Gradient RMS and contrast features. The percentage of cells positively stained for SA-β-gal was determined by the bright-field mean pixel intensity of each cell (lower values corresponds to higher staining).

### Cell cycle analysis

24h following irradiation, NCI-H460 cells were pulsed with 10μM BrdU for 3h at 37°C, and detached and trypsinized attached cells collected and fixed in 4% PFA. After consecutive incubations in Denaturation Solution (2N HCl/Triton X-100 in PBS) and Neutralization Solution (0.1M Na2B4O7, pH 8.5) cells were incubated with Antibody Solution (PBS, 0.5% Tween 20, 1% BSA, 1μg anti-BrdU-FITC conjugated antibody (eBioscience)) and then resuspended in DAPI solution. Samples were analyzed by ImageStream X Mark II as above. At least 2×10^4^ cells were collected from each sample in each biological repeat.

### Protein analysis

Cells were lysed in PLB buffer (10mM NaPO4 pH 7.5, 5mM EDTA, 100mM NaCl, 1% Triton X-100, 1% Na deoxycholate, 0.1% SDS) supplemented with 10μl/ml 0.1M PMSF and 1% protease and phosphatase inhibitor cocktails. For vimentin analysis, cell pellets were lysed in 2% SDS, 50mM Tris-HCl, pH 6.8 or SDS sample buffer (62.5mM Tris-HCl, pH 6.8, 2% SDS, 10% glycerol, 0.1M DTT). Proteins were separated by SDS-PAGE and transferred to nitrocellulose membranes, which were incubated with antibodies against vinculin (Sigma, cat# SA-V9131), cleaved caspase-3 (Cell Signaling, cat# cs9664), tubulin (Sigma, cat# T9026), p21 (Santa Cruz, cat# 6246), GAPDH (EMD Millipore, cat# MAB374), HtrA2 (Cell Signaling, cat# cs2176), vimentin (Sigma, cat #V6389), p53 PAb421 antibody (Bartek et al., 1990). Detection was done with either HRP-conjugated goat anti-mouse or anti-rabbit secondary antibodies (Jackson ImmunoResearch), followed by enhanced chemiluminescence (EZ-ECL, Biological Industries Israel Beit-Haemek Ltd.).

### siRNA functional screening

A custom made siRNA library targeting PCD genes (Dharmacon, Table S4) was reverse transfected into NCI-H460 cells plated in two 96-well plates. Several of the siRNAs were present in both plates and served as internal standards. Non-targeting siRNA NT5 was used as control. 48h after transfection, one set of the siRNA library pair was irradiated, while its pair was left untreated. Cell viability was measured by CellTiter Glo 48h post-irradiation. The experiment was performed in duplicate and repeated 4 times. Each siRNA readout was log transformed, followed by subtraction of the average siNT5 readouts for each plate respectively (irradiated or untreated). The fold-change was obtained by dividing mean values of irradiated by untreated for each siRNA. The statistical difference between irradiated and untreated samples was assessed using a paired T-test, followed by FDR correction (False Discovery Rate, using p.adjust function) comparing four logarithmic values each for untreated cells and irradiated cells. For both tests, a *p*<0.05 was considered significant. All calculations were performed with R.

### Mass spectrometry based proteomic analysis

Mass spectrometry was performed at the G-INCPM unit, Weizmann Institute. Total protein lysates were prepared from non-irradiated and irradiated cells, treated with DMSO or Ucf-101, 48h post-irradiation in four biological replicates. Cells were lysed in buffer (5% SDS, 50 mM Tris-HCl pH 7.4) followed by sonication. Samples were first subjected to in-solution tryptic digestion using the S-Trap method (by Protifi). The resulting peptides were fractionated offline using high pH reversed phase chromatography, followed by online nanoflow liquid chromatography (nanoAcquity) coupled to high resolution, high mass accuracy mass spectrometry (Thermo Q-Exactive HF). Samples from each fraction were analyzed on the instrument separately, and within each fraction, in a random order in discovery mode. Data processing was performed using MaxQuant v1.6.6.0. The data was searched with the Andromeda search engine against the human proteome database appended with common lab protein contaminants and the following modifications: fixed modification (cysteine carbamidomethylation), variable modifications (methionine oxidation), and protein N-terminal acetylation. The quantitative comparisons were calculated using Perseus v1.6.0.7. Decoy hits were filtered out, and only proteins that were detected in at least two replicates of at least one experimental group were kept. A difference ratio for each comparison was calculated based on the average measured protein intensity for each experimental group. A difference in protein level was considered significant if all the following conditions were fulfilled: more than one peptide per protein was identified, the difference ratio was >2 or <0.5, and a *p*-value lower than 0.05 was calculated. Gene Ontology analysis of mass spec results was done using DAVID v6.8 functional annotation analysis (Huang et al., 2009a; Huang et al., 2009b) or IPA (QIAGEN Inc., https://www.qiagenbioinformatics.com/products/ingenuitypathway-analysis) (Krämer et al., 2014). Top canonical pathways were defined as those with (−log(p-value)>3).

### Statistical analysis

Statistical analyses were performed using Excel software or R. Values of *p* < 0.05 were considered significant. Specific tests applied to each set of experiments are described in figure legends. Sample variance equivalence was determined by F-test, where appropriate.

## Acknowledgments

We would like to acknowledge Prof. V. Krizhanovsky (Weizmann Institute) for expertise and insightful discussions.

## Data Availability

The mass spectrometry proteomic datasets produced in this study are available in the PRIDE PXD020972 repository (http://www.ebi.ac.uk/pride/archive/projects/PXD020972) **Username:**; **Password:** GkCQMcNI.

## List of Abbreviations

BrdU: bromodeoxyuridine
CTG: CellTiter-Glo
IF: intermediate filament
KD: knock-down
PCD: programmed cell death
PI: propidium iodide
SA-β-Gal: senescence-associated β-galactosidase activity
SASP: senescence-associated secretory phenotype

